# Whole genome sequencing of field isolates reveals extensive genetic diversity in *Plasmodium vivax* from Colombia

**DOI:** 10.1101/025338

**Authors:** David J. Winter, M. Andreína Pacheco, Andres F. Vallejo, Rachel S. Schwartz, Myriam Arevalo-Herrera, Socrates Herrera, Reed A. Cartwright, Ananias A. Escalante

## Abstract

*Plasmodium vivax* is the most prevalent malarial species in South America and exerts a substantial burden on the populations it affects. The control and eventual elimination of *P. vivax* are global health priorities. Genomic research contributes to this objective by improving our understanding of the biology of *P. vivax* and through the development of new genetic markers that can be used to monitor efforts to reduce malaria transmission.

Here we analyze whole-genome data from eight field samples from a region in Cordóba, Colombia where malaria is endemic. We find considerable genetic diversity within this population, a result that contrasts with earlier studies suggesting that *P. vivax* had limited diversity in the Americas. We also identify a selective sweep around a substitution known to confer resistance to sulphadoxine-pyrimethamine (SP). This is the first observation of a selective sweep for SP resistance in this species. These results indicate that *P. vivax* has been exposed to SP pressure even when the drug is not in use as a first line treatment for patients afflicted by this parasite. We identify multiple non-synonymous substitutions in three other genes known to be involved with drug resistance in *Plasmodium* species. Finally, we found extensive microsatellite polymorphisms. Using this information we developed 18 polymorphic and easy to score microsatellite loci that can be used in epidemiological investigations in South America.

**Author Summary:** Although *P. vivax* is not as deadly as the more widely studied *P. falciparum,* it remains a pressing global health problem. Here we report the results of a whole-genome study of *P. vivax* from Cordóba, Colombia, in South America. This parasite is the most prevalent in this region. We show that the parasite population is genetically diverse, which is contrary to expectations from earlier studies from the Americas. We also find molecular evidence that resistance to an anti-malarial drug has arisen recently in this region. This selective sweep indicates that the parasite has been exposed to a drug that is not used as first-line treatment for this malaria parasite. In addition to extensive single nucleotide and microsatellite polymorphism, we report 18 new genetic loci that might be helpful for fine-scale studies of this species in the Americas.

## Introduction

Despite significant advancements toward malaria control and elimination, about 40% of the world’s population remains at risk of infection by one of the four protozoan species that commonly cause the disease [1]. Among the human malarias, *P. vivax* is the parasite with the most morbidity outside Africa [1]. *P. vivax* differs from the more widely studied *P. falciparum* in aspects of its life cycle, disease severity, geographic distribution, ecology, and evolutionary history [2–7], raising concerns that gaps in our knowledge about its basic biology may compromise its control (see [8] for a detailed discussion).

Genomic approaches provide important tools to study hard-to-culture parasites such as *P. vivax.* For example, genome-wide scans performed on samples from a natural parasite population can identify regions of the genome subject to strong selection. Studies using this approach in *P. falciparum* have contributed to the study of drug resistance and adaptation to the host immune system in that species [9–12]. However, this approach has not yet been widely applied in *P. vivax*.

Previous population genetic studies have found that *P. vivax* populations in many regions of the Americas are less diverse than those from Asia or Oceania [8, 13-17]. Similarly, a recent whole-genome study found limited genetic diversity in a population from the Amazon basin of Peru [18]. It remains unclear whether these results are reflective of populations in the New World generally, the geographical sampling of those particular studies, or the loci sampled in earlier studies. Two recent studies have suggested that *P. vivax* populations in the Americas may harbor more genetic diversity than previously thought. First, a genome-wide comparison revealed substantial genetic divergence among three parasite lineages isolated in the Americas and maintained in non-human primates [19]. Second, recent population studies on the mitochondrial genome have shown high levels of divergence and limited gene flow among populations in the region [20]. This pattern indicates that *P. vivax* populations in the Americas likely have a complex history, with divergent populations harboring differing levels of genetic diversity.

Genomic studies can also support efforts to control and eliminate malaria from a given region by identifying genetic markers that will be informative for fine-scale population genetic studies in that region. Molecular epidemiological investigations rely on multilocus genotyping of SNPs or microsatellites [21] to investigate patterns of population structure and gene flow. Although the high mutation rates of microsatellite loci makes them ideal markers for such studies, only a few loci are currently in use [21]. These existing loci were developed using data from a small number of populations. As a result, some of these loci fail to amplify in samples from other localities [22]. Whole-genome studies offer the opportunity to identify new loci that are both polymorphic in a given region and able to be score reliably (e.g. by selecting loci with simple repeat motifs).In addition to identifying patterns of transmission within a region, these markers can be used to distinguish local cases (the result of remaining malaria transmission) from those that are introduced from another region. Ascertaining the source of the parasites detected in a given case is critical for evaluating the success of interventions during an elimination program.

Here we take a whole-genome approach to characterize the genetic variation of field isolates from single-lineage *P. vivax* infections from Northern Colombia (specifically, Tierralta, Department of Córdoba), an area in South America with seasonal transmission [23]. In addition to assessing the genetic diversity within this population, we examine patterns of diversity across the *P. vivax* genome and find evidence for a recent selective sweep likely associated with resistance to a drug that is not prescribed for treating *P. vivax* malaria. We also develop 18 new microsatellite loci for fine scale studies in Colombia.

## Methods

### Ethics statement

A passive surveillance study was conducted between 2011 and 2013 in outpatient clinics located in Tierralta [23]. The study protocol was approved by the Institutional Review Board (IRB) affiliated to the Malaria Vaccine and Drug Development Center (MVDC, Cali-Colombia). Patients with malaria infections containing more than 5,000 parasites per *μ*L of blood as determined by microscopic examination of Giemsa-stained thick blood smears received oral and written explanations about the study and, after expressing their willingness to participate, were requested to sign an informed consent (IC) previously approved by the Institutional Review Board (IRB) affiliated to the MVDC. IC from each adult individual or from the parents or guardians of children under 18 years of age was obtained. Individuals between 7 and 17 years old were asked to sign an additional informed assent. A trained physician of the study staff completed a standard clinical evaluation and a physical examination in all malaria symptomatic subjects. The local health provider treated individuals as soon as the blood sample had been drawn, using national antimalarial therapy protocol of the Colombian Ministry of Health and Social Protection [23]. Specifically, patients infected with *P. vivax* were treated orally with chloroquine (25 mg per kg provided in three doses) and primaquine (0.25 mg per kg daily for 14 days).

### Sample collection

A brief description of each patients infection status and medical history is provided in S1 Table. We collected 10 mL of blood from each patient and stored each blood sample in EDTA. In order to eliminate as much human DNA as possible, each sample was diluted in one volume of PBS and filtered with a CF11 column (≈ 3*g*) that had previously been rinsed with PBS. A new column was used for each 5 mL of sample. Each filtered sample was centrifuged at 1,000 g for 10 minutes. The supernatant was discarded and the red blood cells (RBC) were kept at -20°C and sent to our laboratory in Cali for processing. The RBCs were suspended in one volume of PBS and aliquoted into 200 *μ*L fractions. DNA was extracted from each aliquot using a PureLink Genomic DNA kit (Invitrogen, USA) following the specifications provided by the manufacturer.

### Sequencing and alignment to reference

Depending on the availability and quality of the sample, we used between 300 ng and 1 mg of DNA to construct sequencing libraries. These libraries were constructed using a Kapa Biosystems DNA Library Preparation Kit (Kapa Biosystems, USA). The resulting fragments were amplified in ten rounds of PCR, using a Kapa HiFi Library Amplification Kit (Kapa Biosystems, USA). Denaturation and clustering were performed using an Illumina cBot. Once the samples were clustered, the flow cell was loaded onto a HiSeq 2000. The run module used was a paired end 2x100 reads. All sequencing and library preparation stages were performed by the DNASU sequencing core at Arizona State University.

We used Bowtie version 2.1.0 [24] to map reads from each sample to a reference genome containing sequences derived from the Salvador I (SalI) strain’s nuclear [25] (build ASM241v1) and apicoplast [26] genomes. The resulting alignments were processed using a modified version of the GATK project’s best practice guidelines [27]. Specifically, we identified and marked potential PCR duplicates using the MarkDuplicates tool from Picard version 1.106 (http://picard.sourceforge.net) and performed local realignment around possible indels using GATK 2.8 [28,29]. Finally, we adjusted the raw base quality scores by running GATK’s BaseRecalibrator tool, treating the set of putative Single Nucleotide Variants (SNVs) identified by SAMtools (0.1.18) [30] as known variants.

To investigate the possibility that our patient samples contain multiple distinct P. *vivax* strains [31,32], we repeated this process using sequencing reads produced from a known single infection. We retrieved reads from the monkey-adapted SalI strain [31] from the NCBI Sequence Read Archive (accession SRS365051) We recorded the frequency of bases matching the reference genome at each site for the patient-derived and single lineage alignments using a custom C++ program (http://dx.doi.org/10.5281/zenodo.18190) that makes use of the BamTools [33] library. For each sample, we also calculated the overall proportion of all sequenced bases that produced a minority allele when mapped to the reference.

### Variant discovery

We called putative SNVs and small indels from the Colombian samples using the GATK UnifiedGenotyper [28]. Because we were able to establish that each patient was infected by a single lineage, we treated samples as haploid. Artifacts produced during the sequencing and mapping of reads to the reference can lead to false positive variant calls [34]. Such calls are more likely in P. vivax due the number of multi-copy gene families and paralogs in this species [35]. In order to account for these potential artifacts, we took a conservative approach to variant calling and removed apparently variant sites that may have resulted from mis-mapped reads. After performing an exploratory analysis comparing properties of our putative variants to a random sample of 100,000 non-variant sites, and using guidelines described by the GATK developers [29], we established the following set of criteria to identify likely false positive variants:

a. Site within 50 kb of a chromosome end (which are dominated by repeats)
b. Average mapping quality phred score <35
c. More than one sample has multiple nucleotides called at the site, such that there are more than two reads that contain minor nucleotides
d. P-value for a Fishers exact test of strand bias <0.001
e. Absolute z-score for Mann-Whitney U-test of mapping quality difference between variant and reference allele containing reads >5
f. Absolute z-score for Mann-Whitney U-test of read-position difference between variant and reference allele containing reads >5
g. Total depth at site ≥ 165x (95th percentile across all sites)

To identify microsatellite loci that are segregating within Colombia and have relatively simple evolutionary histories, we filtered the indels labeled as Short Tandem Repeats by UnifiedGenotyper by removing any matching the following criteria:

a. Locus has non-perfect repeats
b. Repeat motif >8bp
c. Locus is monomorphic among Colombian samples

We produced final variant sets for SNVs and microsatellites by removing all sites that matched at least one of the criteria listed above using PyVCF (http://pyvcf.readthedocs.org/). We produced a functional annotation for each polymorphic SNV with snpeff [36] using the Ensembl functional annotation of the SalI reference (build ASM241v1.23) as input.

### Validation of variants

We validated our SNV calling procedure by running the steps described above on a genome alignment generated from reads previously produced from the SalI strain [31] Because these reads represent an independent sequencing of the same strain that was used to produce the reference genome, we expect very few non-reference alleles. We also tested the effect of including low-coverage samples in our variant calling pipeline by repeating this procedure with the SalI alignment down-sampled to 2x coverage. We further tested the validity of filtered variant sites by searching for apparently singleton SNVs (those found only once in our population sample) in previously reported SNVs from other studies [19,31].

### Oligonucleotide primers for polymorphic microsatellites DNA markers

We designed PCR primers for 18 of the microsatellite loci identified using the above criteria. We first chose a subset of these putative microsatellites that were distributed across different *P. vivax* chromosomes, then developed markers using two strategies. First, nine loci were identified on alignments of conserved regions between the *P. cynomolgi* and *P. vivax* genomes. (SalI *P. vivax* genome data available in NCBI). Second, nine loci were identified on conserved regions between SalI strain and Colombian samples from Tierralta.

All the alignments were made using ClustalX v2.0.12 and Muscle as implemented in SeaView v4.3.5 [37–39]. Dyes Hex and 6-FAM were used for labelling the forward primers. A complete characterization of the 18 microsatellites loci and the primers we used to amplify them is provided in S1 Text and S2 Table.

### Population genetics

We calculated 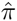 and Watterson’s estimator of the population mutation rate 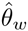 for the whole genome, distinct genomic features (i.e. sequences falling in exons, intron, untranslated regions and intergenic regions), and in 10 kb windows across each chromosome. We accounted for the varying levels of sequence coverage among our samples by using missing-data estimators for these measures [40]. Rather than setting an arbitrary coverage level at which a sample should be considered missing for a given site, we used our data and variant calling approach to identify the number of samples from which we could call a variant if it was present at each site in the genome. We first created new “reference genome” sequences by switching each unambiguous base in the SalI reference following Table 1. We then called variants against these “shifted” genomes using the same procedure described above (including filtering steps). At each site, a sample was considered missing if no variant could be called for that sample using the true reference genome or any of the shifted references.

**Table 1.** Scheme used to produced alternative reference genomes.

|  |  |  |  |  |
| --- | --- | --- | --- | --- |
| Reference | A | C | G | T |
| First shift | T | A | C | G |
| Second shift | G | T | A | C |
| Third shift | C | G | T | A |

We used PyBedtools [41,42] to generate genomic windows, and extract polymorphisms from various sequence classes. We used diversity statistics calculated from genomic windows to identify genomic regions with unusually high or low genetic diversity. Specifically, we identified windows with values in the 1st or 99th percentile of either measure, having first removed windows for which less than 85% of sites were callable. We discovered a region of particularly low diversity surrounding the *dhps* gene, so focused on this gene by calculating each statistic for the set of overlapping windows, each 10 kb wide and 500 bp apart from each other.

## Results

### Genome sequencing

We generated between 18 and 36 million paired-end reads from each sample (Table 2). Obtaining genomic sequences from clinical isolates of *P. vivax* is complicated by the presence of human DNA in parasite-containing blood samples. Although we took steps to remove leukocytes from each of our samples, the proportion of reads that could be mapped to the SalI reference genome differed markedly among samples, ranging from <1%–28%. These differences were reflected in the mean sequencing coverage achieved for each sample, which varies from less than one read per base in sample 500, to greater than 40 reads for sample 499.

**Table 2.**
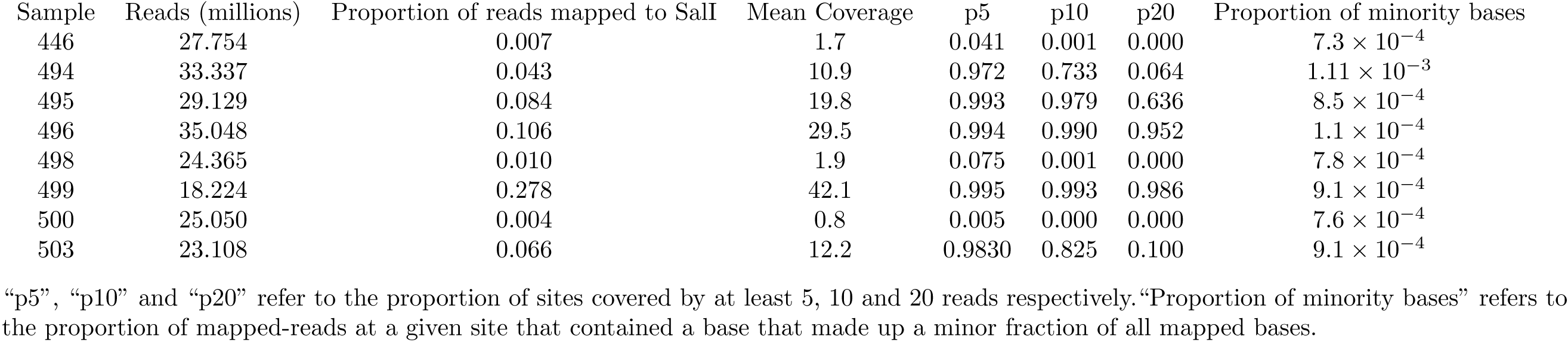
Summary of sequencing data.

Despite the presence of some poorly covered samples, our mapped reads allow comparison between multiple individuals for the vast majority of the *P. vivax* genome (Fig. 1). Because low-coverage samples still provide valuable data for variant calling at some sites and can often be reliably genotyped for those sites known to contain segregating variants, we included all of our patient samples in subsequent analyses.

**Figure 1.**
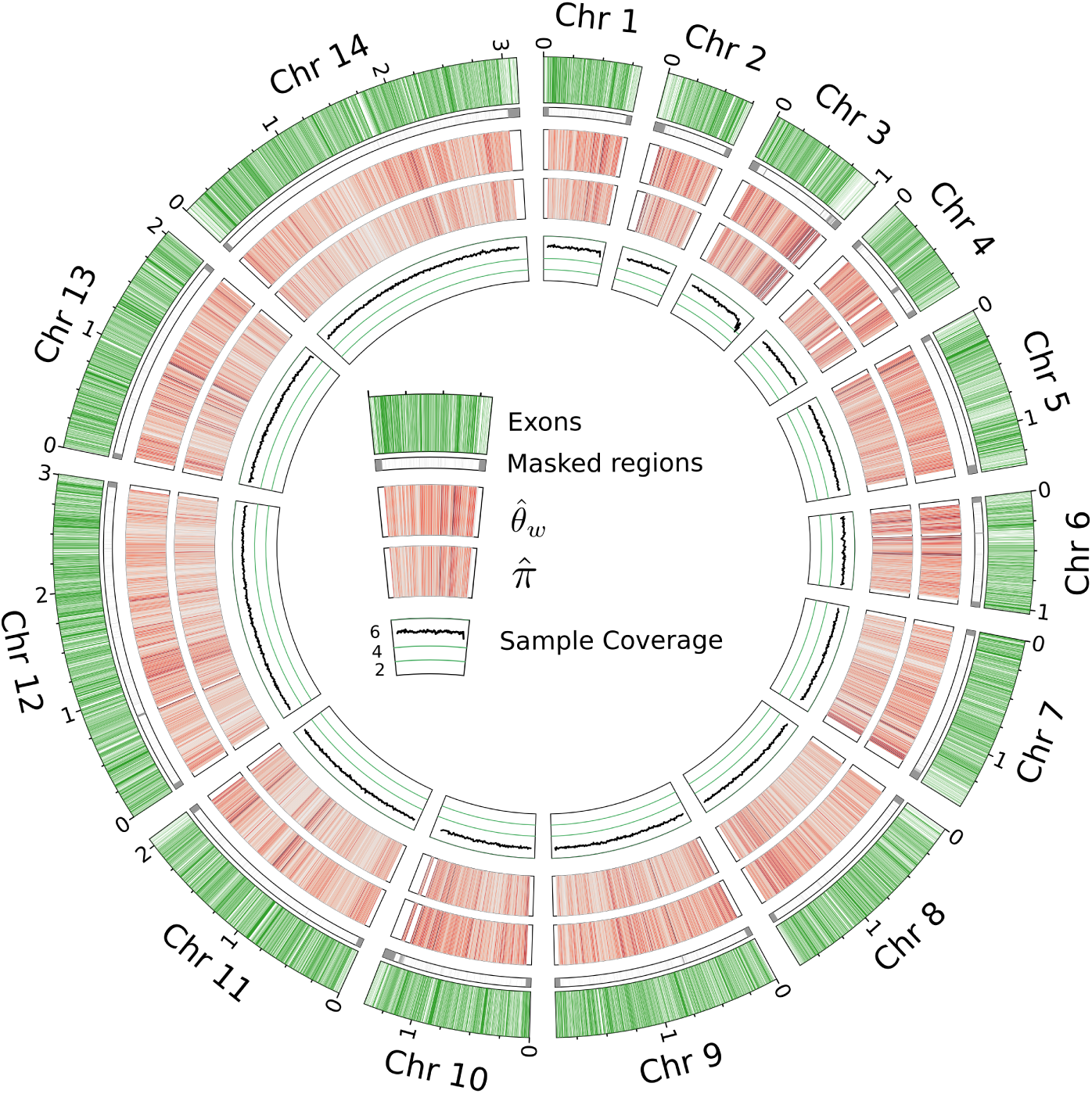
Summary of genomic data: Segments from outside to inside: *P. vivax* chromosomes with exonic regions shaded green; Regions excluded from variant calling in this study; heatmap of 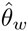 in 10kb windows, high-diversity regions are darker and the maximum value is 0.0018; heatmap of 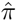 in 10kb windows; Mean number of samples covered per base in 10kb windows

### All sampled patients were infected by only one *P. vivax* strain

The presence of multiple distinct parasite lineages within a single host has presented a barrier to population genetic analysis in previous whole-genome studies of *P. vivax*. Because *Plasmodium* merozoites are haploid in the vertebrate host, the presence of such multiple infections in a patient can be inferred by the presence of multiple alleles at different loci. In the context of high-throughput sequencing, these additional alleles manifest as an excess of bases with intermediate frequencies at sites in a genome alignment. This result contrasts with the distribution expected from singly infected patients, where only rare sequencing errors will produce minority bases [31]. We tested our samples for multiple infections by comparing the distribution of minor base frequencies in our alignments with the same distribution in an alignment produced from a known single infection (Fig. 2). In both the known single infection strain and our samples, minority bases were present only at low frequencies. This pattern contrasts distinctly with the high proportion of intermediate-frequency bases expected from mixed infections [31].

**Figure 2.**
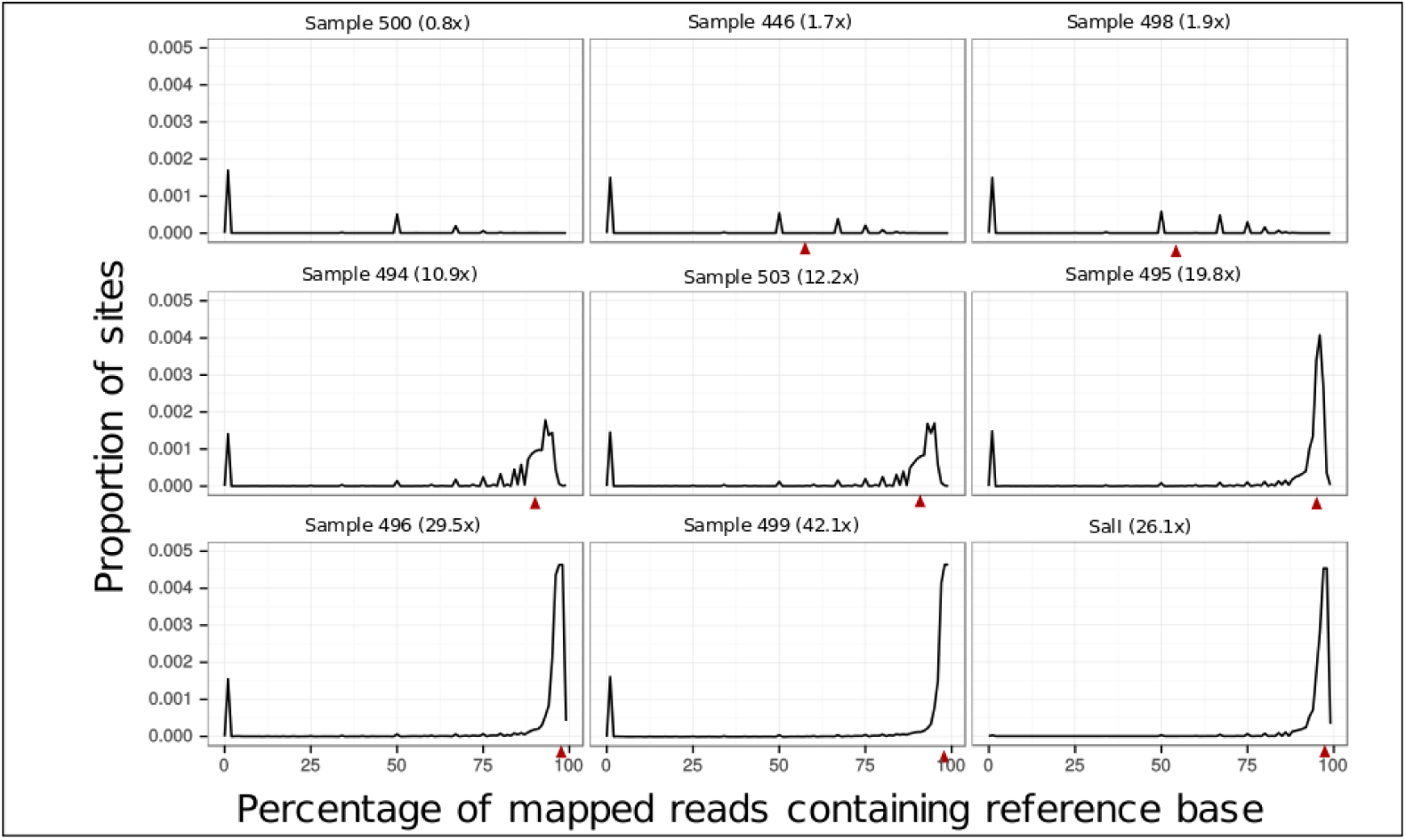
Reference base frequency spectra: The first eight panels represent the distribution of the frequency of reference bases in sequencing reads produced from our samples across all sites in the *P. vivax* genome. In the final panel the same distribution is graphed for reads generated from SalI, a known single infection. Red triangles represent the frequency expected if one non-reference containing read was mapped to a site with average sequencing coverage for that sample (given in parentheses after the sample name). The majority of sites from all samples contain only non-reference bases; these sites were removed to allow clearer visualization of these distributions.

Because the shape of the distribution of minor base frequencies will be less informative for samples with low coverage, we also examined the proportion of all sequenced bases that were in the minority relative to other reads aligned to the same site. Again, these results are similar to those from SalI (Table 2). The proportion of minority bases in reads produced from a known single infection was 7.3 × 10^−4^, while for our data this proportion was between 7.3 × 10^−4^ and 1.11 × 10^−3^. The fact that no samples had the base frequency-distributions characteristic of mixed infections, and that the low-coverage samples did not produce more minor bases than other samples, suggests that all of our samples can be considered single infections.

### Variant discovery

We validated our SNV calling procedure by applying it to a set of sequencing reads produced independently from the same strain used to assemble the reference genome [31]. Using this procedure, we called a total of 34 SNVs from the >22 million base pair reference genome, none of which were called as polymorphic alleles in our patient-derived data. Repeating this procedure with a lower coverage (2x) dataset generated fewer SNVs (18) including only one new putative variant. The small number of variants called from the reference data confirms the conservative nature of our variant calling procedure.

In total, we identified 33,855 non-reference SNV alleles among the Colombian samples, of which 3,594 where fixed difference and 30,261 of which were polymorphic (Table 3). The total number of SNVs we detect is comparable to numbers found in studies of field isolates from other regions. The number of SNVs in our Colombian samples is slightly less than that from samples from Madagascan and Cambodian populations, which contained 41,630 and 45,417 SNVs respectively in the genomic regions included in our study (Fig. 3) [31]; however, it was three times the number identified in the same genomic regions in a recent study using isolates from the Amazon basin in Peru [18]. Approximately two-thirds of the Peruvian SNVs (7,232) are also present in our Colombian samples. However, the Madagascan and Cambodian samples share many alleles that are absent in either South American population (11,618).

**Table 3.** Summary of SNV data.

|  | Total | Exonic | Intronic | 3'UTR | 5'UTR | Intergenic |
| --- | --- | --- | --- | --- | --- | --- |
| Number of SNVs | 33855 | 16413 | 3107 | 808 | 1328 | 14333 |
| Density | 1.8 | 1.2 | 2.4 | 1.9 | 1.8 | 2.0 |
| $\theta_w$ | $7.0 \times 10^{-4}$ | $5.9 \times 10^{-4}$ | $7.9 \times 10^{-4}$ | $6.5 \times 10^{-4}$ | $6.4 \times 10^{-4}$ | $7.5 \times 10^{-4}$ |
| $\pi$ | $6.8 \times 10^{-4}$ | $5.7 \times 10^{-4}$ | $7.5 \times 10^{-4}$ | $6.2 \times 10^{-4}$ | $6.1 \times 10^{-4}$ | $7.2 \times 10^{-4}$ |

**Figure 3.**
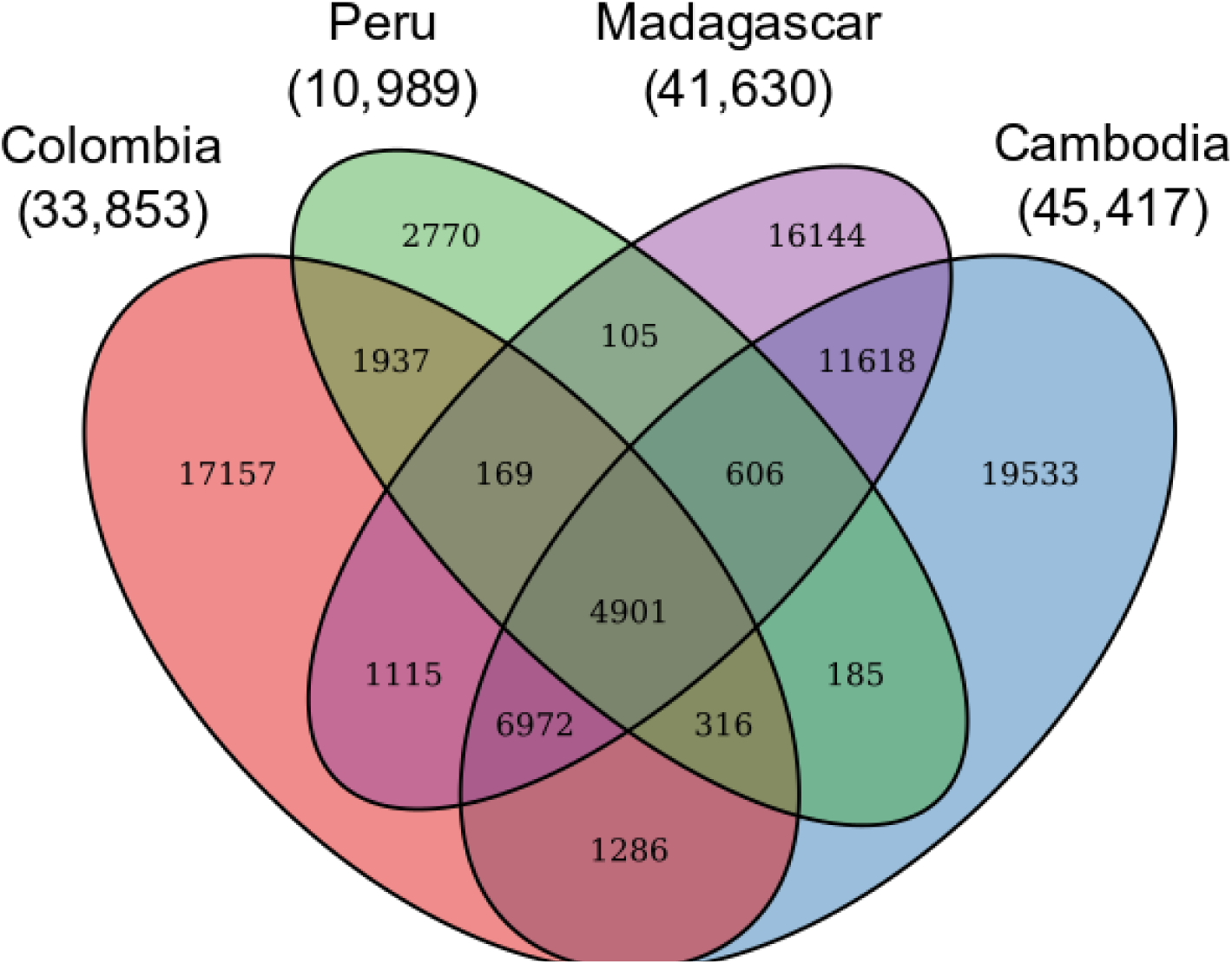
Allele sharing among *P. vivax* populations:. Note: Ellipses for Peruvian [18] Cambodian and Madagascan [31] populations include only those SNVs reported in the cited studies and not included in a region that was excluded in our study.

SNVs are relatively less common in exonic sites (1.2 SNVs per kb) than untranslated, intronic or intergenic sites (1.8-2.4 SNVs per kb) (Table 3). The ratio of non-synonymous to synonymous SNVs in exons is 1:1.51, a result which is lower than the ≈ 1:4 ratio predicted for the *P. vivax* genome under neutrality [31]. This ratio, and the relative densities of SNVs in different sequence classes are very similar to those previously reported from Madagascan and Cambodian populations [31].

Among polymorphic SNVs, 12,913 (42.7%) were recorded in only one sample (Table 4). However, many of these apparent singletons have been recorded in other studies. When we compare our SNVs to a catalogue of previously reported *P. vivax* SNVs [18,19,31] only 6,854 (20.2%) are unique to this study. Our low-coverage samples did not produce more singletons per callable-site than those with higher sequencing coverage (Table 4), suggesting the remaining singletons are not simply artifacts introduced by including these samples.

**Table 4.** Breakdown of singletons.

| Sample | Singletons this study | Singletons all studies | Callable sites |
| --- | --- | --- | --- |
| 446 | 660 (78.7) | 242 (28.9) | 38.1 |
| 498 | 10 (1) | 8 (0.8) | 45.4 |
| 499 | 2974 (158.5) | 1623 (86.5) | 85.3 |
| 494 | 3134 (168.0) | 1606 (86.1) | 84.8 |
| 495 | 837 (44.7) | 520 (27.8) | 85.1 |
| 496 | 2361 (126.0) | 1291 (68.9) | 85.2 |
| 500 | 122 (35.3) | 32 (9.3) | 15.7 |
| 503 | 2806 (150.1) | 1523 (81.4) | 85.0 |
| Total | 12913 (111.9) | 6854 (59.4) | — |

Values in parentheses are number of singleton SNVs per million callable sites.
Values in the Callable sites column are the proportion of all sites that were callable for a given sample.

We identified 789 putative microsatellite loci that met our filtering criteria. To demonstrate the ability of whole-genome studies to develop new markers, we designed PCR primers for 18 of these loci (choosing markers that were well-spread among the 14 *P. vivax* chromosomes). We were able to generate PCR amplicons for each locus, and 16 were shown to be polymorphic within the validation panel, with between 2 and 4 loci segregating in this population (Table 5). The alleles of each locus could be determined easily, as demonstrated by the electropherograms shown in S1 Fig.

**Table 5.** Summary of microsatellite loci discovered in this study.

| Name | Position | Motif | Repeats | Alleles (genomes) | Alleles (validation) |
| --- | --- | --- | --- | --- | --- |
| CLAIM1 | 2:96021 | ATGC | 5–7 | 4 | 4 |
| CLAIM2 | 2:261105 | AC | 6–13 | 4 | 4 |
| CLAIM3 | 5:280204 | AACAGC | 8–11 | 4 | 3 |
| CLAIM4 | 5:1139547 | AT | 8–9 | 3 | 4 |
| CLAIM5* | 5:1195225 | AAAT | 6–10 | 4 | 4 |
| CLAIM6* | 5:1248584 | AT | 8–10 | 3 | 2 |
| CLAIM7 | 6:436676 | AT | 8–9 | 3 | 1 |
| CLAIM8* | 7:761767 | AT | 8–9 | 3 | 2 |
| CLAIM9 | 7:1343057 | AG | 8–12 | 3 | 1 |
| CLAIM10* | 8:439431 | AT | 8–10 | 4 | 4 |
| CLAIM11 | 9:1048573 | AAAT | 6–7 | 3 | 4 |
| CLAIM12* | 9:1454922 | AT | 8–9 | 3 | 2 |
| CLAIM13* | 10:1348254 | AT | 8–9 | 3 | 2 |
| CLAIM14 | 10:1368121 | AT | 3–8 | 3 | 3 |
| CLAIM15 | 12:2076009 | AC | 7–11 | 4 | 3 |
| CLAIM16* | 12:2101147 | AT | 8–9 | 3 | 2 |
| CLAIM17* | 13:950389 | AT | 7–8 | 3 | 2 |
| CLAIM18 | 14:1469652 | AT | 6–14 | 4 | 3 |

Loci marked with an asterisk were developed by comparing *P. vivax* sequences to *P. cynmolyi*. For the numbers of alleles, “genomes” refers to the number identified from genomic data and “validation” to the number detected from the validation panel. PCR primers for each locus are given in S2 Table.

### Population genetics

Because each of our patient samples represents a single parasite lineage, we can use standard population genetic analyses to investigate the evolutionary and demographic processes shaping *P. vivax* genomes in Colombia.

We calculated two measures of genetic diversity: nucleotide diversity 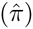 and Watterson’s estimator 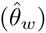 of the population mutation rate [43,44]. Across the whole genome, we estimate 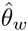 to be 7.0 × 10^−4^. The estimate for nucleotide diversity is slightly lower at 6.8 × 10^−4^. For both diversity measures, genetic diversity is lowest in exonic regions, then increasingly higher in 3′ and 5′ untranslated regions of transcripts, intergenic regions, and introns (Table 3).

### A selective sweep around *dhps*

We identified regions of the genome with unusually high or low genetic diversity in this population (S3 Table). The most striking result of this analysis is an extended region of homozygosity on chromosome 14, which includes a 10kb window with no polymorphic SNVs despite having a mean of 6.35 samples contributing data. When we narrowed our focus to this region by calculating 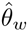 in overlapping windows, we found that the Dihydropteroate Synthetase gene *(dhps*) was at the center of this region of low diversity (Fig. 4). Although there are no polymorphisms within this region, all eight of our samples contain a non-reference allele resulting from a G to C substitution in the second exon of *dhps*. The substitution is non-synonymous and leads to the A383G amino acid substitution that has been associated with sulphadoxine resistance in numerous previous studies of *P. vivax* [45]. This pattern of low diversity surrounding a fixed substitution is the classic sign of a hard selective sweep [46,47], in which a single mutant or migrant allele is rapidly fixed by selection. No other non-reference alleles were present in the *dhps* gene.

**Figure 4.**
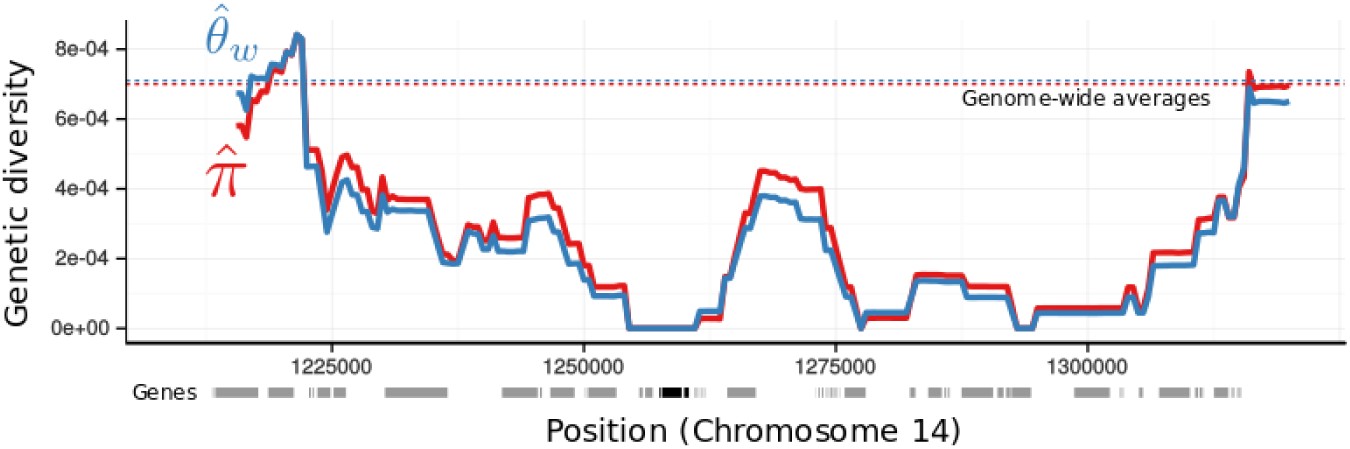
A selective sweep around *dhps*: Two measures of genetic diversity (y-axis), were calculated in overlapping windows across a portion of Chromosome 14 (positions in x-axis). The dashed path under the x-axis represents the position of exons in the current annotation of the *P. vivax* reference genome. Exons of *dhps* are shaded black, all others are shaded gray. Dashed lines represent the genome-wide average of each diversity measure.

Most of the remaining genes that overlap with other high- or low-diversity windows encode proteins for which there is no functional annotation. Nevertheless, the high-diversity windows include antigen and surface protein genes such as *msp7* and *vir* family proteins, which are known to be under balancing selection in *P. vivax* [19]. Because our conservative variant calling approach removed difficult-to-align multi-copy gene families, it is likely we excluded other genes under balancing selection.

We also examined other genes thought to be involved in drug resistance in *P. vivax* 6. Alleles of the dihydrofolate reductase *(dhfr*) gene are known to confer resistance to pyrimethamine, a drug administrated together with sulphadoxine (SP). We did not observe evidence of a selective sweep at the *dhfr* locus. However, all samples for which a genotype could be called reliably (six of eight) have non-synonymous SNVs leading to both S58R and S117N amino acid substitutions. Both of these substitutions have been associated with SP resistance in *P. vivax* [45]. There are two distinct nucleotide variants encoding the S58R substitutions in our population samples, with three parasites having AGC>AGA mutations in the 58th codon, and three others having an AGC>CGC mutation. We found a total of 13 additional non-synonymous variants in the ATP-cassette binding proteins *(pvmdr1* and PVX_124085, a homolog of *P. falciparum mrp* proteins). There were no such variants in GTP cyclohydrase, Chloroquine resistance transporter ortholog, or Kelch 13, which are all considered possible drug resistance genes [18].

## Discussion

### Genetic diversity in Colombia

The genetic diversity estimated from our Colombian *P. vivax* population is similar to, though slightly lower than, estimates derived from a global sample of *P. falciparum* (where 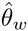 has been estimated to be 1.03 × 10^−3^ using isolates from Africa, America, Asia and Oceania [48]). Although it is well known that *P. vivax* is more genetically diverse than *P. falciparum* globally, this finding is important as it demonstrates that *P. vivax* control programs face genetically diverse populations even in relatively small spatial scales in South America.

It is difficult to compare the genetic diversity of our sample to that of other *P. vivax* populations. Most studies that report estimates of genetic diversity for this species are focused on clinically important epitopes or a few markers. On the other hand, whole-genome studies cannot usually report diversity statistics due to multiplicity of infection in their samples. However, we can compare our heterozygosity estimates to those produced from one large scale population genetic study.

A study of using 5.6 kb of non-coding DNA from *P. vivax* isolates from across India [49] reported 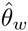 values ranging (from 1.3 × 10^−3^ − 3 × 10^−3^). These values are somewhat higher than our diversity estimates from introns 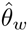 or intergenic regions 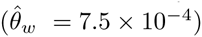 but within the range of values we calculate from 10kb windows.

We can also make a crude comparison between our results and those from other whole-genome studies [18, 31] by comparing the number of non-reference alleles found in each population. Despite our relatively small number of well-covered samples, restricted geographic range, and the conservative approach to variant calling, we detected 33,855 SNVs. When we apply the same masking criteria used in this study to the variants reported from an unknown number of parasite lineages from Cambodia and Madagascar we arrive at comparable number of SNVs (41,630 and 45,417 respectively). Our study discovered considerably more variants than a recent study of *P. vivax* isolates from the Amazon basin in Peru (10,989 variants). It is likely that this difference reflects the low genetic diversity of the particular Peruvian population studied. Simply counting non-reference alleles does not provide a direct comparison of genetic diversity between populations, especially when the number of parasite lineages sampled in other studies is not known. Nevertheless, these results demonstrate the Colombian population sampled here does not contain an unusually low number of non-reference alleles as might be expected from a population with low genetic diversity.

Comparing the whole-genome studies also highlights the degree of allele-sharing among populations (Fig. 3). Approximately two-thirds of the SNVs detected from a *P. vivax* population in Peru were also detected in our Colombian samples, and 11,618 alleles are shared by the Cambodian and Madagascan populations but absent from both South American samples. This pattern may represent genetic differentiation between New World and Old World populations, although it is important to note that differences between these studies may also reflect different variant calling procedures. A complete understanding of the global structure of *P. vivax* populations, and the relative diversity of populations on different continents, will require many more population samples and a consistent approach to variant calling.

Together these findings differ substantially from the perception that South American *P. vivax* populations in general have low diversity as a result of a simple evolutionary history [19,20]. Some populations, including the Peruvian population discussed above, do indeed have low genetic diversity, which may be the result of recent local introductions or expansions from a few founders following malaria control programs [50]. On the other hand, the Colombian population studied here has substantial genetic diversity. It has been suggested that such diversity could be result of complex demographic processes involving multiple introductions and admixture among lineages in broad temporal and spatial scales [20]. A comprehensive population genomic study of South America would be required to understand the extent of such genetic polymorphism and the processes involved in its maintenance. Nevertheless, our results demonstrate that South American *P. vivax* populations do not universally have low genetic diversity.

Our heterozygosity estimates are calculated under the assumption that each sampled patient was infected by a single parasite lineage. If multiple *P. vivax* lineages were present in some patient samples our estimates may be inflated. However, we do not believe this is likely. None of our samples show the excess of intermediate frequency bases expected to arise from sequencing of mixed infections (Fig. 2). Furthermore, our lowest coverage samples, which would be most likely to generate extra variant calls if multiple infections were present, do not produce an excess of rare bases or singletons SNVs. The apparent lack of multiple infections among our patient samples may reflect local conditions in Córdoba, including seasonal transmission of the disease and the widespread use of primaquine, a drug that eliminates dormant infections. Indeed, three of our patients are known to have received primaquine in the 12 months prior to providing their sample.

### Signatures of natural selection

We identified genomic windows with exceptionally high or low genetic diversity. High genetic diversity may be maintained by balancing selection, while regions of low diversity may be subject to strong purifying selection or be the product of recent selective sweeps. The majority of genes contained within these windows encode proteins for which little is known. However, some of the high diversity windows overlap with antigen and surface protein genes that are thought to be subject to balancing selection in *P. vivax* globally [19]. It is possible that genes in other windows of exceptional diversity have likewise been subject to natural selection; this could be confirmed with a larger population sample and thus a statistically more powerful genome scan.

We also looked at patterns of diversity more broadly by comparing results from different genomic features. All measures of diversity were lowest in exon sequences, followed by untranslated regions, intergenic spaces, and introns. This pattern, along with the relative lack of non-synonymous SNVs, is consistent with earlier studies demonstrating that purifying selection has a strong effect on protein coding genes in *P. vivax* [51]. It is interesting to note that our estimates of genetic diversity were higher for introns than intergenic regions. This pattern has been reported in *P. falciparum* [19]; it may reflect the presence of conserved, but unannotated, genes in what are currently considered intergenic regions [48].

### Drug resistance alleles in Colombia

Our analysis of low diversity regions revealed a selective sweep associated with the A383G allele of *dhps,* which has previously been associated with resistance to Sulfadoxine [45]. Although resistance to SP treatment, and indeed resistance mediated by this particular mutation, is a well known phenomenon, this result is interesting for two reasons. First, by identifying a selective sweep around this mutation we are able to demonstrate that SP resistance has arisen within Colombia via the rapid fixation of a single allele. This finding, combined with the fact that another *dhps* allele (A385G) is most commonly associated with SP resistance in Madagascar [52], French Guiana [52], India [53], Iran [54], Pakistan [55], Thailand [56] and China [57], suggests SP resistance has come about through multiple independent origins. Similar repeated evolution of *dhps* resistant mutations has been reported in *P. falciparum* [58–61]

The evolution of SP resistance is also interesting in an operational context because it demonstrates that this *P. vivax* population has been subject to drug pressure from SP, a drug that has not been part of the approved treatment for uncomplicated *P. vivax* malaria in Colombia (where the drugs of choice are still chloroquine-primaquine combination therapy). SP has been used to treat *P. falciparum* infections in Colombia, so it is possible this selective pressure has arisen from misdiagnosis of *P. vivax* infections or the use of SP to treat mixed *P. vivax*-*P. falciparum* infections. It is also possible that poor compliance with national drug policies, including self-medication by some patients with access to antifolates or the long half-life of antifolate drugs could lead to *P. vivax* infections coming into contact with SP.

The region surrounding the *dhfr* does not show the pattern of decreased genetic diversity associated with a hard selective sweep. In this case, all genotyped samples have two SP-resistance alleles (S58R and S117N), with the first allele encoded by two distinct SNVs. Thus, SP-resistance alleles in each gene have somewhat different histories, with *dhps* A383G entering the population once and being rapidly driven toward fixation, but *dhfr* resistance arising from two separate alleles that have been maintained in the population. This pattern differs from the one found in *P. falciparum,* where *dhfr* mutations associated with drug resistance are fixed as the result of a selective sweep, whereas *dhps* mutations are still segregating with sensitive alleles in the population [59,61].

We detected non-synonymous variants in two other genes thought to be involved with drug resistance in *Plasmodium* species. There are eight non-synonymous variants in PVX_124085. The phenotypic effects of these variants are not known, but the changes to the *P. falciparum* ortholog of this gene have been associated with decreased sensitivity of primaquine [62] and antifolate drugs [63]. This study is the third time that an excess of non-synonymous mutations in this gene has been recorded from a population in South America [18,64]; the clinical significance of this repeated finding should be investigated.

We identified five non-synonymous mutations in *pvmdr1,* three of which are known from populations in Asia and Madagascar [52,65]. Mutations in the *P. falciparum* ortholog of this gene are associated with decreased sensitivity to chloroquine, and *pvmdr1* variants are thus considered putative chloroquine-resistance alleles. Among the variants we report, only Y976F has been associated with decreased sensitivity to chloroquine *(in vitro* and with a modest effect size) [66]. There is little evidence for drug failure with chloroquine in South America at present. Nevertheless, the presence of these alleles in a South American population warrants further investigation and presents an opportunity to test for an association between *pvmdr1* alleles and sensitivity to the drug in a clinical setting.

We also compared the variants we report from putative drug resistance genes with those reported from another South American population in the Amazon basin of Peru (Table 6). These populations share many alleles, including both SP resistane alleles in *dhfr* and five amino acid substitutions in PVX_124085. In contrast, four of the five variants we reported from *pvmdr1* are not present in the Peruvian population and there are no non-synonymous substitutions in Peruvian *dhps* sequences. These results demonstrate the importance of local information in designing control programs, as each population contains distinct drug resistance alleles that may generate distinct responses to different treatments. In addition, the fact that two populations separated by a considerable geographical distance, as well as by the Andes, share multiple alleles that are identical by state at the nucleotide level, suggests that it is possible for drug resistance alleles to spread by gene flow between distant populations in South America.

**Table 6.** Non-synonynous variants at drug resistance loci.

| Locus | Chromosome | Position | Codon | Amino Acid | Number of samples |
| --- | --- | --- | --- | --- | --- |
| *dhps* | 14 | 1257856 | gCc/gGc | A383G | 8 (8) |
| *dhfr* | 5 | 964761 | Agc/Cgc | S58R* | 4 (7) |
|  | 5 | 964763 | agC/agA* | S58R* | 3 (7) |
|  | 5 | 964939 | aGc/aAc | S117N* | 6 (6) |
| *pvmdr1* | 10 | 363169 | tAc/tTc | Y976F | 3 (8) |
|  | 10 | 363223 | aCg/aTg* | T958M* | 8 (8) |
|  | 10 | 363374 | Atg/Ctg | M908L | 8 (8) |
|  | 10 | 364598 | Gat/Aat | D500N | 4 (8) |
|  | 10 | 365435 | Gtg/Ttg | V221L | 5 (8) |
| PVX_124085 | 14 | 2043859 | Caa/Gaa* | Q1419E* | 1 (7) |
|  | 14 | 2044528 | Ccg/Tcg | P1196S | 3 (8) |
|  | 14 | 2044798 | Gcg/Tcg* | A1106S* | 2 (6) |
|  | 14 | 2044975 | Ctg/Atg | L1047M | 1 (7) |
|  | 14 | 2045050 | Gtg/Atg* | V1022M* | 7 (7) |
|  | 14 | 2046072 | aGc/aTc | S681I | 3 (6) |
|  | 14 | 2047233 | aGg/aTg* | R294M* | 7 (7) |
|  | 14 | 2047816 | Gaa/Caa* | E100Q* | 2 (6) |

“Number of samples” refers to the number of genomic sequences that contained this allele in the present study, the number of samples from which any allele could be reliably called appears in parentheses. Variants marked with an asterisk were also recorded from a population in the Amazon basin in Peru [18]. Note the numbering of amino acid substitutions in PVX\_124085 differs between these studies as a result of the different annotations used in each case.

### Microsatellite loci for *P. vivax* studies in the Americas

Although population genomic studies offer a unique view into the biology of *P.vivax,* smaller-scale studies that use genotypes from only a few loci will remain important in malaria research. One important result from this study is a set of new microsatellite loci that can be used in fine scale population genetic and molecular epidemiological studies in Colombia. Microsatellite loci are particularly useful for such studies, as their relatively high mutation rates can generate highly polymorphic loci. As a result, population genetic signals in microsatellite loci can reflect demographic events occurring at short time scales, including epidemiological events [50, 67]. These markers can also be used for population assignment and in testing for multiple infections [21,22,68,69].

Thus far, 160 microsatellites have been found in the genome of *P. vivax* [22]; however, many of these loci fail to amplify in some populations [21,22]. It is not surprising that loci developed in one region are not necessarily informative in others: microsatellites have complex evolutionary histories [70] and high potential for homoplasy [71]. Thus, the widespread application of microsatellite loci to epidemiological problems will require the development of new markers known to amplify and be polymorphic within specific populations. We found 789 putatively polymorphic microsatellite loci from our whole-genome sequencing, demonstrating that the 160 markers currently used in *P. vivax* represent only a small proportion of loci available in this species. Moreover, we demonstrated that the loci we detected in our whole-genome sequencing can be developed into useful makers. The 18 markers we developed yield patterns of repeats that are easy to score in populations in the Pacific Coast of Colombia and show high levels of polymorphism. Whether these markers will be useful at broader geographic scales remains to be seen, but the specific markers we developed will be useful for fine-scale studies in this region, where malaria elimination is currently being considered.

## Conclusion

Our results add to growing evidence that P. vivax populations are genetically diverse. Even at a small spatial scale in Colombia, this *P. vivax* population harbors levels of genetic diversity similar to a global sample of *P. falciparum.* The diversity we observed appears to be inconsistant with demographic scenarios in which South American *P. vivax* populations were derived from a recent introduction, possibly associated with a severe population bottleneck [17,72]. As such, our study lends further support to the idea that *P. vivax* populations in the Americas have a more complex history than was previously thought [20].

Our study also demonstrates that genomic studies of natural populations of *P. vivax* can provide insights into how parasite populations react to control strategies. Specifically, we identified a selective sweep associated with resistance for SP, a drug that is not used to treat *P. vivax* in Colombia. This resistance indicates a possible spillover effect form of a drug that is primarily used to treat *P. falciparum*. The operational consequences of these results require additional investigations. Future studies with additional samples may detect additional regions under selection and thus contribute to the identification of vaccine targets [73] or other clinically relevant phenotypes. Finally, we used our genomic data to develop a set of microsatellite markers that are both easy to genotype and known to be polymorphic within this population. These markers will aid future epidemiological studies and our understanding of malaria transmission and demography in Colombia.

## Supporting Information Legends

### S1 Text

Methods used for microsatellite development

**S1 Table.** Patient summary information.

**S2 Table.** Characterization of 18 polymorphic *P. vivax* microsatellite loci. Size ranges of PCR products (in base pairs) are given for six of Colombian *P. vivax* isolates. SalI was used as a positive control. Fluorescent dyes (Hex and 6-FAM) were used to label forward primers only. ML: Motif length and No.A: allele numbers.

**S3 Table.** Extreme diversity windows. 10kb windows with unusually high or low genetic diversity. Values for 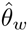 and 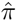 are × 10^−3^. The values in the “Start” column are the position of the start of the genomic window in kb.

**S1 Fig.** Electropherograms showing peak profiles for 18 polymorphic microsatellite loci. The y-axis corresponds to fluorescence intensity (arbitrary units) and the x-axis is the PCR product length in base pairs (bp). The amplitude of the each peak in base pairs (bp) is shown in boxes underneath the peaks. The range of allele sizes for these small datasets is also given for each locus.

## Acknowledgments

We are grateful to Elizabeth Winzeler and David Serre, who provided raw SNP data from their own studies of *P. vivax* populations. We also thank Scott Roy, Marcelo Ferreira and two anonymous reviewers for their comments on this paper.

